# Popularizing recombinant baculovirus-derived OneBac system for scaling-up production of all recombinant adeno-associated virus vector serotypes

**DOI:** 10.1101/2020.11.01.363606

**Authors:** Yang Wu, Zengpeng Han, Mingzhu Duan, Liangyu Jiang, Tiantian Tian, Dingyu Jin, Qitian Wang, Fuqiang Xu

## Abstract

Recombinant adeno-associated virus (rAAV) has been widely used as an efficient transgenic vector in biomedical research, as well as gene therapy. Serotype-associated transduction efficiency, tissue- or cell-type tropism and immunological profile are major considerations in the various applications of rAAVs. There are increasing needs for different serotypes of rAAV, either naturally isolated or artificially engineered. However, affordable and scalable production of a desired serotype of rAAV remains very difficult, especially for researchers lacking relevant experience. On the basis of our previously established single recombinant baculovirus expression vector (BEV)-derived OneBac system, we have optimized the process and expanded the rAAV production range to the full range of serotypes rAAV1-13. Firstly, the AAV *Cap* gene was optimized to translate by ribosome leaky scanning and the gene of interest (GOI) was cloned into the pFD/Cap-(ITR-GOI)-Rep2 shutte plasmid. Following the classical Bac-to-Bac method, sufficient BEV stock containing all rAAV packaging elements can be quickly obtained. Finally, we can repeatedly scale up production of rAAVs in one week by using a single BEV to infect suspension-cultured Sf9 cells. The rAAV1-13 show relatively high yields ranging from 5×10^4^ to 4×10^5^ VG/cell. More than 1×10^15^ VG purified rAAVs can be easily obtained from 5 L suspension-cultured Sf9 cells. As expected, rAAV serotypes 1-13 show different potencies for *in vitro* transduction and cell-type tropisms. In summary, the single BEV-derived OneBac system should prove popular for laboratory scaling-up production of any serotype of rAAV.

## Introduction

Recombinant adeno-associated virus (rAAV), a safe and efficient transgenic vector, has been widely used in basic and clinical research in biomedicine, as well as gene therapy [1, 2]. rAAVs play important roles especially for its excellent *in-vivo* performance in cutting-edge areas of biological research, such as neuroscience and CRISPR/Cas9-mediated gene editing [3, 4]. Since 2012, when the first rAAV-based gene therapy drug Glybera (rAAV1-LPL vector) was approved, another two rAAV drugs Luxturna (rAAV2-hRPE65 vector) and Zolgensma (rAAV9-SMN vector) have been successively approved in 2017 and 2019, respectively [5]. So far, according to the gene therapy clinical trial report website www.abedia.com/wiley, there are more than 250 ongoing rAAV-based gene therapy clinical trials worldwide, including for hemophilia A and B, Duchenne muscular dystrophy (DMD), and Parkinson’s disease.

rAAV was first developed as a gene transfer vector in mammalian cells from wild-type AAV in 1984 [6]. AAV is a small non-enveloped virus belonging to the *Parvoviridae* family, *Dependovirus* genus, and contains a linear single-stranded DNA genome approximately 4.7 kb in length [2]. To date, 13 different AAV serotypes (AAV1 to AAV13) have been identified based on the phylogenetic classification, and more than 100 variants from human and primate origin are known. Variants differ in their capsid structures and display variable transduction efficiency, tissue- or cell-type tropism, and immunological profile [7-9]. Efforts have been made to isolate or engineer new AAV serotypes [10-13]. Some artificially engineered AAV serotypes have been created and became very popular owing to their excellent features, such as the AAV-DJ [14], AAV-Anc80L65 [15] and AAV-PHP.B [16]. Meanwhile, exploring and clarifying the features of different AAV serotypes is also very important for the applications of rAAVs [17-19].

The rAAV genome contains a gene of interest (GOI) flanked by the AAV inverted terminal repeats (ITR), which is packaged in rAAV icosahedral capsid about 20-25 nm in diameter. The AAV ITRs serve as *cis* elements, while the AAV *Rep, Cap* and helper genes are provided in *trans* for viral replication and packaging [20]. Currently, the classical helper-free system using triple plasmid cotransfection of HEK293 cells is widely used for the small-scale laboratory manufacturing of rAAVs [21]. The three plasmids are pAAV-RC, pAAV-(ITR-GOI) and pAAV-helper, respectively. Although this method has been optimized for production of rAAV at large scale, optimization with suspension-cultured HEK293 cell line is complex, hard to popularize and expensive [22, 23]. The proper expression levels of *Rep* and *Cap* genes related to the pAAV-RC construct are crucial for generating high-titer rAAV [19]. Due to the gene transcriptional regulations varying among different AAV serotypes, it is not easy to design a proper pAAV-RC construct for a new rAAV serotype [24-26]. Moreover, the yields and quality of rAAV could vary drastically from batch to batch due to the amount and ratio of plasmids and transfection reagents used, cell viability, and operating mode [27, 28]. Due to existing safety risks, adenovirus (ADV)- or herpes simplex virus (HSV)-infected mammalian cell-mediated rAAV production systems are not widely used [29]. Therefore, affordable and scalable production of rAAVs of a desired serotype remains very difficult for most common laboratories.

The development of baculovirus expression vector (BEV)-mediated rAAV production systems (Bac systems) were originally aimed to facilitate large-scale and low-cost production of rAAV. In 2002, Urabe et al. first reported the production of rAAV upon infecting insect Sf9 cells with three BEVs: BEV/Cap, BEV/Rep and BEV/(ITR-GOI) [30]. In order to improve the genetic instability of BEV/Rep and the low co-infection ratio, the optimized *Cap and Rep* genes expression cassettes were integrated in a dual-functional BEV/Cap-Rep with two different strategies: artificial introns used by Chen in 2008 [31] or ribosome leaky scanning used by Smith et al. in 2009 [32]. However, due to intrinsic low co-infection ratio of the earlier multi-BEV infection-based ThreeBac and TwoBac systems, rAAV yields rarely exceed 5×10^4^ vector genomes (VG) per cell by this method. In 2009, Aslanidi et al. established a Sf9/Cap-Rep packaging cell line-dependent OneBac system, which overcome the drawback of the low co-infection ratio. The rAAV was produced upon BEV/(ITR-GOI) infecting the inducible Sf9/Cap-Rep packaging cell line harboring silent copies of AAV *Rep* and *Cap* genes. Yields exceed 10^5^ VG/cell by this method, which is about 10-fold higher than previous Bac systems [33]. In 2014, Mietzsch et al. further expanded this OneBac system to produce the serotypes rAAV1-12 [34]. In 2019, we reported an improved Sf9/GFP-Rep packaging cell line dependent OneBac system, which is more flexible to switching between different serotypes of *Cap* genes using simple BEV/Cap-(ITR-GOI) reconstruction [35]. However, the difficulty to obtain such high yield packaging cell lines seriously reduced their potential popularization for laboratory use.

We also developed a single BEV/Cap-(ITR-GOI)-Rep-derived packaging cell line-independent OneBac system in 2018 [36], which might be a better choice for laboratory production of rAAVs. In this study, we further optimized the process and expanded the rAAV production range to serotype 1-13 based on the new OneBac system. We can quickly construct the required shuttle plasmid pFD/Cap-(ITR-GOI)-Rep2 and integrate all rAAV packaging elements by conventional molecular cloning operations, which only needs the AAV *Cap* gene and GOI-expressing casette to be changed according to the required serotype. Then, the BEV/Cap-(ITR-GOI)-Rep2 can be generated according to the classical Bac-to-Bac system protocol. After that, we can repeatedly scale up production of rAAVs just in one week by using a single BEV to infect suspension-cultured Sf9 cells. All the rAAVs 1-13 show relatively high yields ranging from 5×10^4^ to 4×10^5^ VG/cell and show different potencies for in vitro transduction and cell-type tropisms. The single BEV-derived OneBac system would be an affordable choice for laboratory scaling-up production of rAAVs of any serotype.

## Materials and Methods

### Plasmid construction

The *Cap* genes of AAV1-13 serotypes were synthesized and/or cloned into pBlueScript II SK(+) plasmids between *HindIII* and *SpeI* cut sites, named as pDonor-Cap1 to pDonor-Cap13. The *Cap* genes of AAV1-13 serotypes are referred to AAV1 (GenBank: NC_002077.1), AAV2 (GenBank: NC_001401.2), AAV3 (GenBank: U48704.1), AAV4 (GenBank: U89790.1), AAV5 (GenBank: NC_006152.1), AAV6 (GenBank: AF028704.1), AAV7 (GenBank: AF513851.1), AAV8 (GenBank: NC_006261.1), AAV9 (GenBank: AY530579.1), AAV10 (GenBank: AY631965.1), AAV11 (GenBank: AY631966.1), AAV12 (GenBank: DQ813647.1) and AAV13 (GenBank: EU285562.1).

To construct the EF1α-GFP-WPRE-pA expression cassette, the EGFP gene was amplified with GFP-F and GFP-R primers, and then cloned into the *BamHI* and *EcoRI* cut sites of the plasmid pAAV-EF1α-Dio-GFP-WPRE-pA (addgene: 37085). The EF1α-GFP-WPRE-pA expression cassette was cut with *NotI*, and then inserted into pFD/Cap2-(ITR-CMV-GFP-pA)-Rep2 as previously described [36] to obtain the pFD/Cap2-(ITR-EF1α-GFP-WPRE-pA)-Rep2 shuttle plasmid.

Then, we used CapX-Pac1-F and CapX-Nhe-R primers to amplify the *CapX* genes, which were then inserted into the pFD/Cap2-(ITR-EF1α-GFP-WPRE-pA)-Rep2, to substitute the *Cap2* gene to yield pFD/CapX-(ITR-EF1α-GFP-WPRE-pA)-Rep2 plasmids. The *CapX* in this study refers to *Cap1* to *Cap13*. The primers are supplied in Supplementary Table S1. The BEVs were generated according to the Bac-to-Bac Baculovirus Expression System protocol (Invitrogen).

### Cell culture

For mammalian cells, human embryonic kidney cell line HEK293T, human cervical cancer epithelial cell line HeLa, human microglia cell line HMC3, human glioma cell lines U87 and U251 were all cultured in Dulbecco’s modified Eagle’s medium (DMEM, Gibco) supplemented with 10% (v/v) fetal bovine serum (FBS), 100 units/ml penicillin, and 100 µg/ml streptomycin at 37 °C, 5% CO2.

For insect cells, Sf9 cells were adherently cultured in Grace’s Insect Medium (Gibco) supplemented with 10% FBS in plates at 28°C, or suspension cultured in Sf-900II SFM (Gibco) in shake flasks at 28°C, 130 rpm.

### BEV production

The BEVs were generated according to the Bac-to-Bac system protocol (Invitrogen). Briefly, the pFD/Cap-(ITR-EF1α-GFP-WPRE-pA)-Rep2 shuttle plasmids were transformed into the DH10Bac *E. coli* strain, and integrated into the baculovirus *Autographa californica* anucleopolyhedrovirus (AcMNPV) genome through Tn7-mediated recombination. About 2 μg positive Bacmid DNA were transfected in 6 well plate cultured Sf9 cells using the Cellfection II reagent. The transfected Sf9 cells showed apparent GFP expression and cytopathic effect after 3-5 days post transfection. The culture supernatants of the transfected Sf9 cells were collected as BEV stock P1. To expand the BEV, the BEV P1 can be used to infect naïve Sf9 cells. Three days post-infection, the culture supernatants were collected as BEV stock P2.

### rAAV production and purification

The Sf9 cells suspension cultured in flask were adjusted to 3×10^6^ cells/ml density with fresh culture medium and infected with BEV/Cap-(ITR-EF1α-GFP-WPRE-pA)-Rep2 at a MOI of 3. Three days post-infection, the infected Sf9 cells were harvested by centrifugation and storage at −80°C until using for downstream purification as described previously [37, 38].

In brief, the cell pellets were suspended in lysis buffer (50 mM Tris, 2 mM MgCl_2_, pH 7.5) and then subjected to three freeze-thaw cycles between liquid nitrogen and 37°C water bath. The crude lysates were added with 50 U/ml benzonase (Sigma) and 150 mM NaCl and incubated at 37°C for 1 hr, then clarified by centrifugation at 2,500×g for 15 min and filtered. The rAAV-containing supernatant was used for virus titration or further purified by iodixanol gradient ultracentrifugation. The iodixanol step density gradients (15%, 25%, 40% and 58%) were prepared by diluting 60% iodixanol (OptiPrep, Sigma) with PBS-MK buffer (1×PBS, 1 mM MgCl_2_, 2.5 mM KCl) and then were successively underlaid into a 39 ml QuickSeal tube (Beckman Coulter). The rAAV-containing supernatant was then gently overlaid onto the gradient and then centrifugated for 2 hr in a 70 Ti rotor at 63,000 rpm,18°C. The fraction obtained from the 40% phase was dialyzed against PBS buffer and then ultrafiltered and concentrated with Amicon Ultra-15 centrifugal filter units (MWCO, 100 kDa, Millipore). The purified rAAVs were stored at −80°C.

### Virus titration

Viruses were titered by quantitative PCR (qPCR) assay using the iQ SYBR Green Supermix kit (Bio-Rad). Standard curves were obtained by 10-fold serial diluting with standard plasmids from 10^8^ to 10^3^ copies per μl. The BEV and rAAV samples were treated as described previously [34]. The quantitative primers were Q-Bac-F and Q-Bac-R for BEV, and Q-WPRE-F and Q-WPRE-R for rAAV, supplied in Supplementary Table S1.

### Western blotting and silver staining

The cells samples were treated and separated using 10% SDS-PAGE electrophoresis. The AAV Rep, Cap and β-tubulin proteins were detected with the anti-Rep monoclonal antibody (clone 303.9, Progen), anti-AAV VP1/VP2/VP3 monoclonal antibody (clone B1, Progen) and monoclonal antibody (66240-1-lg, Proteintech), respectively.

The purified rAAV samples were separated using 10% SDS-PAGE electrophoresis. The gels were then silver stained with a Fast Silver Stain Kit (Beyotime) according to the manufacturer’s protocol.

### Transmission electron microscopy

Transmission electron microscopy was performed on each purified rAAV1-13. About 5 μl rAAV sample was dropped onto a 400-mesh carbon-coated copper grid. After 2 min incubation, the unabsorbed liquid was blotted off the grid at the edge with Whatman’s filter paper. The wet grid was negative stained with 2% phosphotungstic acid (pH 7.4) for 2 min. Then, the grid was washed with a drop of ddH2O and air-dired. Finally, grids were examined on a Hitachi H-7000A transmission electron microscope. Negative stained images were obtained at 100000×.

### In vitro transduction assay

HEK293T, HeLa, HMC3, U87 and U251 cells were seeded in 96-well plates about 8×10^3^ cells/well. When the cells reached 70% confluence, the medium was replaced by DMEM-2% FBS. Then, the 5-fold serial diluted samples of rAAVs were added into 96-well plates to infect these cells. After 2 days post-infection, the GFP fluorescence was observed by fluorescence microscopy (Olympus X71).

## Results

### Generation of the BEVs for producing rAAV1-13

We have developed a novel BEV integrated with all packaging elements for the production of rAAV as previously described [36]. To facilitate virus testing, a new expression cassette containing an EGFP reporter gene driven by EF1α promoter was generated through the *Not1* cut sites of the pFD/Cap2-(ITR-CMV-GFP-pA)-Rep2 shuttle plasmid. For convenience, the codon-optimized *Rep2* gene was used for producing different serotypes rAAV. In addition to *Cap2* gene, the other serotype *Cap* genes modified for ribosome leaky scanning were cloned into the pFD/Cap2-(ITR-EF1α-GFP-WPRE-pA)-Rep2 shuttle plasmid through the *Pac1* and *Nhe1* cut sites to substitute the *Cap2* gene. The start codons of AAV Capsid proteins (VP1,VP2 and VP3) were changed to CTG, ACG and ATG, respectively. After cloning, the pFD/Cap-(ITR-EF1α-GFP-WPRE-pA)-Rep2 plasmid integrated all rAAV packaging elements were prepared for each serotype of rAAV (Fig.1). Then, the shuttle plasmids for each of the 13 serotypes were separately transformed into the DH10Bac strains. After Tn7 transposon-mediated recombination, the positive Bacmid DNA was obtained and then transfected into Sf9 cells. After three to five days post-transfection, Bacmid DNA-transfected Sf9 cells for each of the 13 serotypes displayed clear GFP fluorescence and cytopathic effect (CPE), (Fig.2A). We collected the culture supernatants of the transfected Sf9 cells with obvious CPE as BEV passage 1 (P1). To expand the BEV stocks, we used the BEV P1 to infect suspension cultured Sf9 cells. After 72 hr post-infection, all of the infected Sf9 cells showed apparent GFP expression and CPE, similar to the Sf9 cells post-transfection (Fig. 2A). We collected the culture supernatants and harvested the cells. The culture supernatants of the BEV P1 infected Sf9 cells were collected as BEV P2.

**Figure 1.**
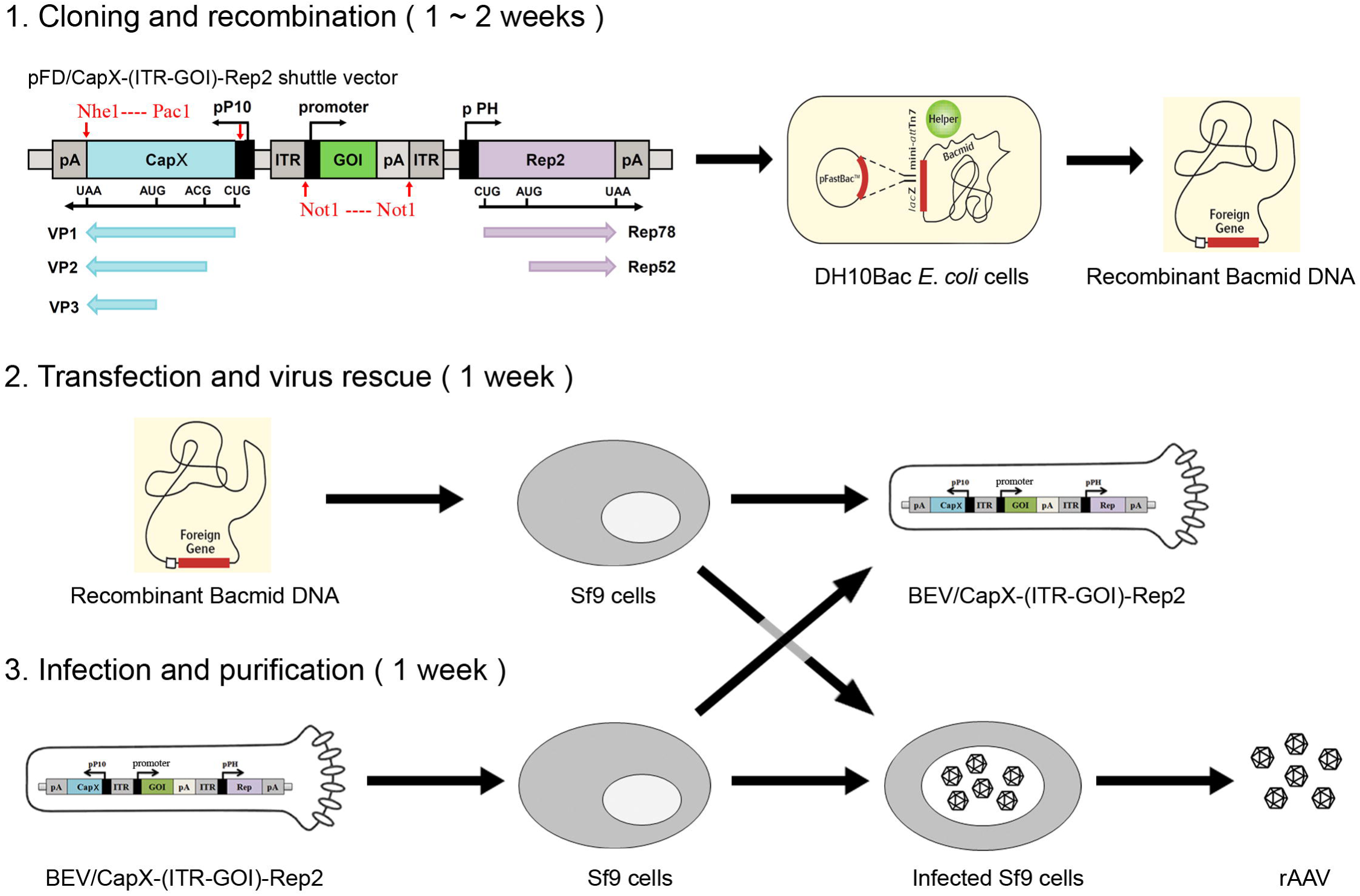
Process schematic for the production of rAAVs by using the novel OneBac system. Step 1: Cloning and recombination. To obtain the required shuttle plasmds, *Cap* genes of the desired serotype, termed as *CapX*, or the expression cassette of the GOI can be quickly cloned into the pFD/CapX-(ITR-GOI)-Rep2 shuttle plasmid through the *PacI/NheI* cut sites or the *NotI* cut sites. Then the pFD/CapX-(ITR-GOI)-Rep2 plasmid was transformed into the DH10Bac *E. coli* strain for Tn7 transposon-mediated recombination. The positive Bacmid DNA containing the rAAV packaging elements were extracted from the transformed DH10Bac strain. Step 1 requires one to two weeks. Step 2: Transfection and virus rescue. The positive Bacmid DNA was used to transfect adherent-cultured Sf9 cells to produce BEV P1 in the cultural supernatant. Step 2 requires one week. Step 3: Infection and purification. The BEV P1 was used to infect suspension-cultured Sf9 cells to expand BEV P2, and for producing the rAAV. After three days post-infection, the infected Sf9 cells were harvested and used for purification of the rAAV. Step 3 requires one week.

**Figure 2.**
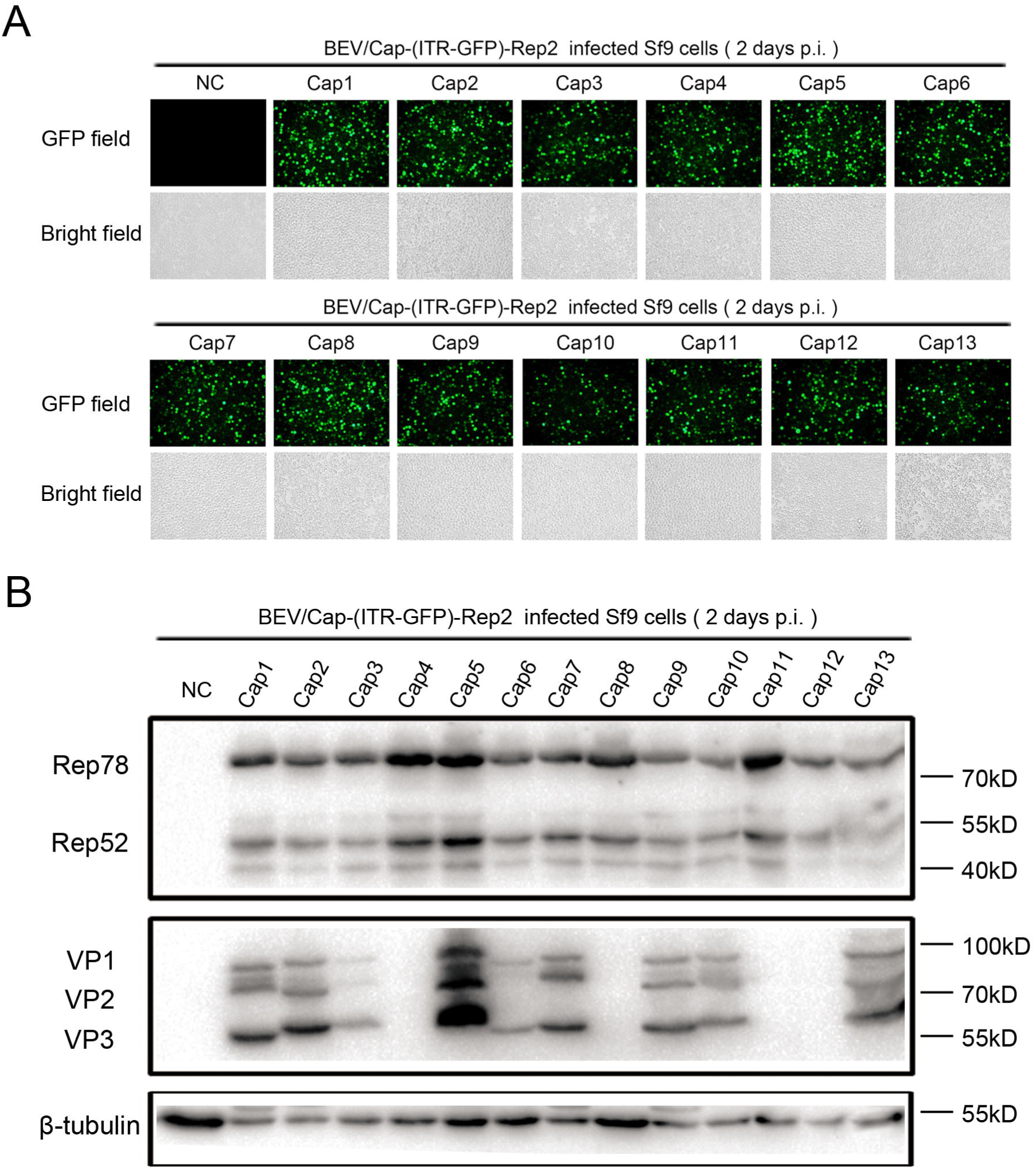
Analysis of gene expression in Sf9 cells during the generation of the BEVs containing the rAAV packaging elements of rAAV serotypes 1-13. A. Fluorescence microscopy observation of GFP expression in Bacmid DNA-transfected Sf9 cells. Approximately 2 μg Bacmid DNA containing rAAV packaging elements of each AAV serotypes were separately transfected into Sf9 cells cultured in 6 well-plates. After three to five days post-transfection, the transfected Sf9 cells were detected by fluorescence microscopy. B. Western blot analysis of Rep and Cap proteins expression in Sf9 cells infected with the BEVs. Suspension-cultured Sf9 cells were infected with BEVs at an MOI of 3. After 2 days post-infection, the infected Sf9 cells were harvested and subjected to western blot analysis. Rep proteins were detected with mAb 303.9. Cap proteins were detected with mAb B1, which could detect all three capsid proteins of most AAV serotypes with the exception of AAV4, 8, 11 and 12. The endogenous cellular protein β-tubulin were detected as internal reference.

In order to detect the different serotype of AAV *Rep* and *Cap* gene expression in the BEV-infected Sf9 cells, the BEV P1 infected Sf9 cells were lysed and analyzed by western blot (Fig. 2B). The Rep78/Rep52 proteins showed comparable levels among different simples as detected by the anti-Rep monoclonal antibody mAb 303.9 (Fig. 2B, upper panel). This indicated that the *Rep2* gene is well expressed in Sf9 cells infected by different BEVs. The expression of different serotype of *Cap* genes were detected by anti-AAV VP1/VP2/VP3 monoclonal antibody mAb B1. As expected for the antibody detection range among the 13 serotypes of Cap proteins, the Cap1, 2, 3, 5, 6, 7, 9, 10, 13 were detected, but the Cap4, 8, 11, 12 could not be detected. For most serotypes, AAV *Cap* genes were revealed as three bands with the correct molecular weight corresponding to the VP1, VP2 and VP3 range from 55kDa to 100kDa. We also noticed that the AAV *Cap5* gene expression levels was obviously higher than the others, while the *Cap3* and *Cap6* genes expression levels were a little lower (Fig. 2B, lower panel). Taken together, the data indicate that BEV/Cap-(ITR-EF1α-GFP-WPRE-pA)-Rep2 were successfully generated for all 13 serotypes.

### Characterization of the rAAV1-13 derived from the BEVs infected Sf9 cells

These BEVs were further used to infect suspension cultured Sf9 cells according to our previously optimized conditions [36]. After three days post-infection, the cells were harvested for purification of rAAVs. Yields ranged from 5×10^4^ to 4×10^5^ VG/cell, (Fig. 3A) with rAAV2, rAAV8 and rAAV9 all exceeding 1×10^5^ VG/cell, which is consistent with previous studies. The Sf9 cells are suitable for scaling-up suspension culture from several microliters to liters in flasks or bio-reactors. In common laboratory conditions, we can easily produce more than 1×10^15^ VG purified rAAVs from 5 L suspension-cultured Sf9 cells in an economical way.

**Figure 3.**
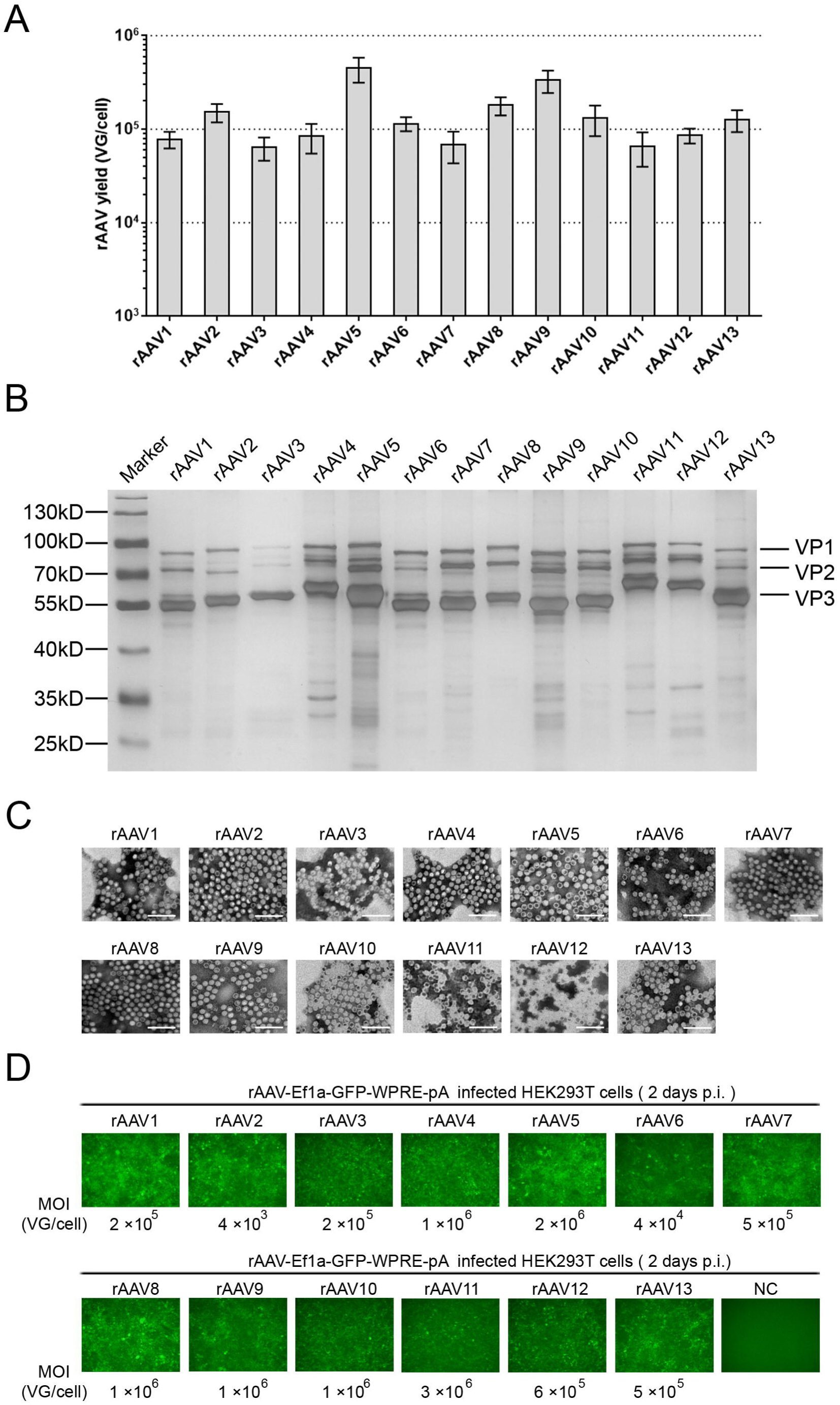
Characterization of the OneBac system-derived rAAV serotypes 1-13. A. The yields of rAAV1-13 derived from the BEV-infected cells, determined as vector genomes per cell (VG/cell). All experiments were performed in triplicate. Mean ± SD values are presented. B. Silver stain of purified rAAV1-13. The purified rAAVs were separated by 10% SDS-PAGE, and then they were analyzed by sliver staining. About 1×10^10^ VG rAAVs were loaded per lane. C. Negative staining transmission electron microscopy analysis of purified rAAV1-13. Representative fields are shown. Original magnification × 100,000. Scale bars, 100 nm. D. Transduction activity analysis of the rAAV1-13. HEK293T cells were infected with gradient diluents of rAAV1-13 at different MOIs and observed under a fluorescence microscope 2 days post-infection. Representative fields are shown.

To characterize the purified rAAV1-13 obtained from the novel OneBac system, we analyzed their biophysical properties and in vitro transduction activity. The rAAVs’ purity was analyzed by silver staining, with all of the purified rAAV1-13 showing very high purity (Fig. 3B). On SDS-PAGE, three bands corresponding to AAV capsid proteins (VP1, VP2 and VP3) were clearly seen within the molecular weight range from 55kDa to 100kDa. The rAAV1-13 Cap proteins (VP1, VP2 and VP3) ratios were expressed at near the expected 1:1:10 ratio with little deviation between different serotypes of rAAV. The morphology of the rAAVs was interrogated by transmission electron microscopy. All serotypes showed similar structures as uniform icosahedral particles with a diameter of 20-25 nm (Fig. 3C). The full, genome-containing particles were white, while the empty particles were dark due to the electron-dense central region of the capsid. The ratios of full particles to empty particles vary among different serotypes of rAAVs. Most serotypes of rAAV show quite high ratios of full particles, whereas rAAV5 is a slightly lower.

Next, the rAAVs’ transduction activity was analysed by infecting HEK293T cells. After 2 days post-infection, obvious GFP fluorescence was seen in all rAAV1-13 infected HEK293T cells with appropriate MOIs (Fig. 3D). The results indicate that all rAAV1-13 had transduction activity, while varying from each other towards HEK293T cells. The results also provide proper initial MOI ranges of 13 serotypes for the rAAV transduction activity assay upon infection with HEK293T cells. In summary, the rAAV1-13 can successfully produced from novel OneBac system in relative high yields and high transduction activity.

### Analysis of the neurotropic properties of the representative serotypes of rAAVs

Due to the new OneBac system being convenient for us to produce rAAVs of different serotypes, we are able to compare between and choose the best serotype for our specific research needs. In the central nervous system, glial cells are the most abundant cell type and are involved in central homeostatic processes, neurodevelopment, immune response and many neurological disorders [39]. rAAVs are a popular transgenic vector for glial cells [40]. Therefore, we analysed the transduction activity of six representative serotypes (rAAV1, 2, 5, 6, 9 and 13) with two commonly used cell lines (HEK293T, HeLa) and three glial cell lines (HMC3, U87 and U251). For each serotype, the five cell lines were infected with identical amounts of rAAVs in 5-fold serial dilutions. After 2 days post-infection, GFP fluorescence was detected by fluorescence microscope. For HEK293T and HeLa cells: rAAV1 and rAAV2 showed comparable transduction activity (Fig. 4A, B); rAAV5, rAAV9 and rAAV13 showed higher transduction activity for HEK293T cells than HeLa cells (Fig. 4C, E, F); rAAV6 showed lower transduction activity for HEK293T cells than HeLa cells (Fig. 4D). For HMC3, U87 and U251 cells: rAAV5 showed comparable transduction activity (Fig. 4C); rAAV1 showed higher transduction activity for HMC3 cell than U87 and U251 cells (Fig. 4A); rAAV2, rAAV6, rAAV9 and rAAV13 show higher transduction activity for U87 cells than HMC3 and U251 cells (Fig. 4B, D, E, F). These results indicate that the transduction activity of different rAAV serotypes varies substantially with different cell types.

**Figure 4.**
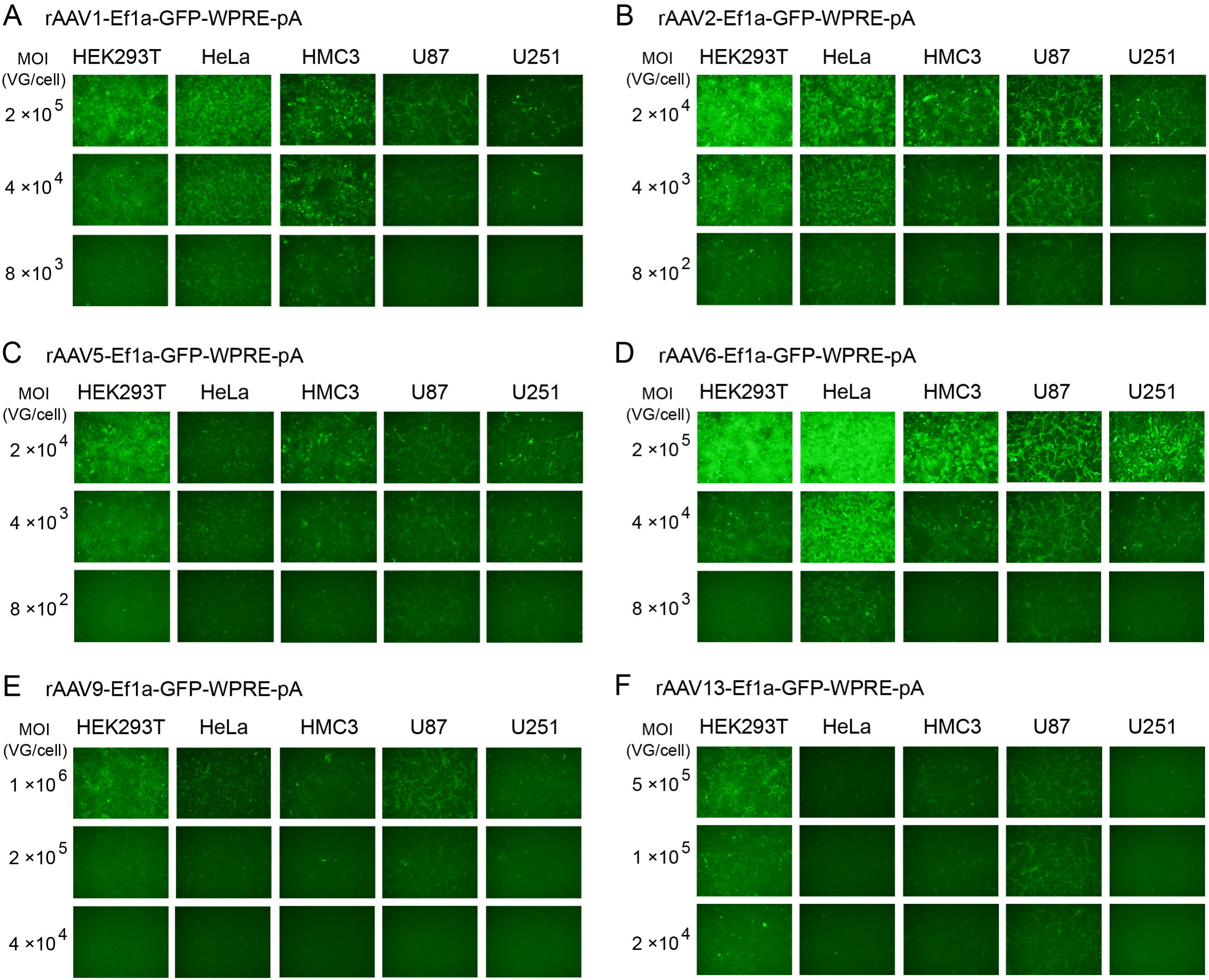
Analysis of the *in vitro* transduction activity of typical serotypes of rAAV derived from the OneBac system using HEK293T, HeLa, HMC3, U87 and U251 cell lines. Cell lines were seeded in 96-well plates at 70% confluence. We infected the cell lines with 5-fold serial dilutions of rAAVs at specified starting MOIs, and then determined transduction activity 2 days post-infection by fluorescent microscopy. The rAAV genome was Ef1a-GFP-WPRE-pA. (A) rAAV1, starting MOI 2×10^5^ VG/cell. (B) rAAV2, starting MOI 2×10^4^ VG/cell. (C) rAAV5, starting MOI 2×10^4^ VG/cell. (D) rAAV6, starting MOI 2×10^5^ VG/cell. (E) rAAV9, starting MOI 1×10^6^ VG/cell. (F) rAAV13, starting MOI 5×10^5^ VG/cell. Representative fields are shown.

## Discussion

In the last decade, the remarkable successes in rAAV-based gene therapy have attracted much attention and investment from big pharmaceutical companies. Given recent industrial and technological advances applied to the processes of large-scale rAAV production, the cost of rAAV drugs will come down to an affordable level for patients in the future. Meanwhile, there are increasing needs for various serotypes of rAAVs in preclinical research with different kinds of animal models, which usually have a dose range from 1×10^11^ VG to 1×10^15^ VG. However, laboratorial rAAV production processes are usually time-consuming, difficult to scale up and lead to batch-to-batch instability of rAAV. Obviously, it is still difficult for researchers to find a proper method that could meet their various demands for rAAV production.

The recent improvements on flexibility and versatility of the OneBac systems have gradually attracted many researchers’ attention. We previously developed a single BEV/Cap-(ITR-GOI)-Rep-based packaging cell line-independent OneBac system. The yields of rAAV2, rAAV8 and rAAV9 all exceeded 1×10^5^ VG/cells. Moreover, the BEV is stable for four serial passages without serious reduction of rAAV yield level [36]. In order to exploit the potential for the production of various rAAVs in the laboratory by using the new OneBac system, we have optimized the process and expanded the rAAV production range to the full range of serotypes rAAV1-13.

The whole process can be divided into three steps (Fig. 1). The first step is cloning and recombination on the *E. coli* strain level, which requires one to two weeks. It is very simple and flexible to switching between different serotypes of *Cap* genes and GOI expression cassettes in the pFD/Cap2-(ITR-GOI)-Rep2 shuttle plasmid. The *Cap* genes modified with ribosome leaky scanning worked very well for the production of most rAAV serotypes (Fig. 2A). The codon-optimized *Rep2* gene also expressed very well for different BEVs at relatively consistent levels (Fig. 2B). The *Rep2* gene is sufficient for the production of different serotypes of rAAVs in insect cells similar as it used in mammalian cells [24]. Then, the shuttle plasmid was transformed into DH10Bac strain for high efficiency Tn7 transposon-mediated recombination with the BEV genome. After a rapid and efficient blue-white screening, the positive recombinant bacteria harboring the Bacmid DNA can be easily obtained. The second step is the transfection and virus rescue on the Sf9 cell level, which takes about one week. The Bacmid DNA was transfected into Sf9 cells to produce the original BEV seed stock termed as BEV Passage 1 (P1). Due to the amount of BEV P1 being small and not sufficient for large scale production of rAAV, we usually have to expand BEV stocks by using the BEV P1 to infect naïve Sf9 cells. The third step is infection and purification, which also takes about one week. The BEV stock was used to infect suspension-cultured Sf9 cells. After 72 h post-infection, the infected Sf9 cells were harvested and used for downstream purification. A variety of purification options are available for the rAAV purification, which can be finished within two to three days. In summary, once sufficient BEV/Cap-(ITR-GOI)-Rep2 stocks are obtained in the initial two to three weeks, we can repeatedly scale up production of various serotypes of rAAVs just in one week by using a single BEV to infect suspension-cultured Sf9 cells without complex process optimization.

In order to speed up the transition from transfection to production, in 2017, Scholz et al. reported the titerless infected-cells preservation and scale-up (TIPS) method that can eliminate repeated virus amplification steps [41]. It could further reduce the time from Bacmid DNA to rAAV products, while reducing the risk of BEV decay. In 2013, Guo et al. developed a simplified rAAV purification method by PEG/(NH_4_)_2_SO_4_ aqueous two-phase partitioning, which is independent with ultrahigh speed gradient centrifugation or chromatography [42]. This approach is a good choice for small-scale rAAV purification in the absence of expensive experiment apparatus. In 2020, Yu et al. reported a method using three-phase partitioning combined with density gradient ultracentrifugation, which is able to purify in excess of 3×10^15^ VG rAAV with just two 38.3 ml Beckman SW28 centrifuge tubes [43]. This method would be an economical and universal choice to purify a large quantity of rAAVs.

The novel OneBac system is suitable for scaling-up production of rAAVs. There are many recombinant proteins, including approved vaccines and therapies, that have been successfully produced in BEV-infected insect cells [44]. The insect Sf9 cells can be scaled up for culture at high cell density in serum-free suspension medium without huge efforts to optimize growth conditions. We make a conservative estimate that 100 ml BEV P2 can be obtained from 2 ml BEV P1 at 50-fold volume expansion by this method. By this method, the yield of the major rAAV serotypes can reach a relatively high level of 5×10^4^ to 4×10^5^ VG/cells (Fig. 3A), which is similar to the high yield level of current Bac systems. The BEV stock P2 is sufficient to infect 5 L suspension-cultured Sf9 cells, which can produce more than 1×10^15^ VG purified rAAV. In consideration of the BEV/Cap-(ITR-GOI)-Rep is stable at least for P4 [36], the rAAV productivity of the new OneBac system could satisfy the demands for most rAAV-based preclinical research.

It is commonly believed that both insect cell- and mammalian cell-based rAAV production systems yield comparable biological activity [34, 45, 46]. Many factors can affect the activity of rAAV during its production. The high rAAV purity can result in serotype- and tissue-independent enhancement of transduction efficiency [47]. Apart from the downstream purification, the upstream packaging methods could yield rAAV which differ in viral capsid protein ratio, virion integrity, and post-translational modifications (PTMs), etc. In 2015, on the basis of the Sf9/Rep-Cap packaging cell-dependent OneBac system, Mietzsch et al. applied a splicing-based strategy to enhance VP1 expression level of rAAV5, which resulted in a 100-fold increase of rAAV5 infectivity [48]. This result is consistent with that the VP1 carrying phospholipase A2 domain in its N-terminus is essential for AAV infectivity [49]. Furthermore, they also found that minimal encapsidation of foreign DNA were seen for rAAV1, rAAV2, rAAV5 and rAAV8 [47, 48, 50]. In 2017, Kondratov et al. further utilized an attenuated Kozak sequence and a leaky ribosome scanning to regulate AAV capsid protein stoichiometry in serotype-specific manner. As a result, the rAAV5 derived from the modified OneBac system showed markedly higher biological potencies than the rAAV5 derived from the HEK293 cells [51]. In 2017, Savy et al found that replacement of truncated ITRs with wild-type and additional AAV2 sequences could improve the rAAV integrity ratio (from 10% to 40%) and reduce non-rAAV encapsidated DNA (up to 10-fold) for rAAV production using the TwoBac system [52]. The PTMs of rAAV capsid proteins differ among different serotypes [53]. A recent study revealed that the rAAV capsid PTMs and rAAV genome methylation differ between two main platforms corresponding to the plasmid-transfected human HEK293 cells and the BEV-infected insect Sf9 cells. It also showed that rAAVs derived from the HEK293 cells are more potent than rAAVs derived from Sf9 cells [54]. However, this study only focused on rAAV1 and rAAV8, and did not illustrate the relevance between these differences and the activity of rAAVs. Therefore, further studies would help to improve the performance of rAAVs produced by Bac systems.

Compared to the complicated *Cap* and *Rep* gene transcriptional regulation designs for the triple-plasmid transfection HEK293 cell-based system, the *Cap* and *R*ep gene constructs using ribosome leaky scanning strategy with the new OneBac system is more simple and worked very well. The majority of rAAV1-13 Rep and Cap protein expression levels were comparable (Fig. 2B). The majority of rAAV1-13 capsid proteins (VP1:VP2:VP3) were expressed close to the prototypic 1:1:10 ratio (Fig. 3B). However, one limitation was that the Cap5 expression level was significantly higher than other serotypes (Fig. 2B), which may have also given rise to lower full particle ratio (Fig. 3C). Further studies should be conducted to optimize the production of different serotypes of rAAVs using the new OneBac system. It is important to choose the best serotype of rAAVs for different applications. rAAV1-13 derived from the new OneBac system showed relatively high yield levels ranging from 5×10^4^ to 4×10^5^ VG/cells, and showed different transduction activity toward HEK293T cells (Fig. 3). The representative serotypes of rAAVs derived from the new OneBac system indeed vary significantly for their cell type tropism (Fig. 4). The convenient and affordable rAAV production method based on the new OneBac system would facilitate the characterization and application of various serotypes of rAAVs. In conclusion, the single BEV/Cap-(ITR-GOI)-Rep2 derived packaging cell line-independent OneBac system will be popular for laboratory scaling-up production of various rAAV serotypes.

## Author Contributions

Y. W. designed and performed the experiments, analyzed the data, and wrote and edited the manuscript. Z.H., M.D. and L.J. performed the experiments and acquired the data. T.T., D.J. and Q.W. carried out viral vector production. F.X. supervised the project and edited the manuscript.

### Acknowledgments

This work was supported financially by the National Natural Science Foundation of China (31500868 to Y.W. and 31771156 to F.X.), Strategic Priority Research Program (B) (XDB32030202 to F.X.), the State Key Program of National Natural Science Foundation of China (31830035 to F.X.) and the Key-Area Research and Development Program of Guangdong Province (2018B030331001 to F.X.).

