## Supplemental Table 1.pdf for "Popularizing recombinant baculovirus-derived OneBac system for scaling-up production of all recombinant adeno-associated virus vector serotypes"

**Supplemental Table S1. Primers for cloning and Q-PCR.**

| Primer | 5'-sequence-3' |
|---|---|
| Cap1-Pac1-F | TGCCTTAATTAAAATACTATACTGTAAATTACATTTTATTTACAATCACTCGACGAAGACTTGATCACCCGGGCCGCCCTGGCTGCCGACGGTTATCTACCCGATTGGCTC |
| Cap3-Pac1-F | TGCCTTAATTAAAATACTATACTGTAAATTACATTTTATTTACAATCACTCGACGAAGACTTGATCACCCGGGCCGCCCTGGCTGCTGACGGTTATCTACCCGATTGGCTC |
| Cap4-Pac1-F | TGCCTTAATTAAAATACTATACTGTAAATTACATTTTATTTACAATCACTCGACGAAGACTTGATCACCCGGGCCGCCCTGGCTGACGGTTACCTTCCAGACTGGCTAGAG |
| Cap5-Pac1-F | TGCCTTAATTAAAATACTATACTGTAAATTACATTTTATTTACAATCACTCGACGAAGACTTGATCACCCGGGCCGCCCTGGCTTTTGTTGATCACCCACCCGATTGGTTG |
| Cap6-Pac1-F | The same as Cap1-Pac1-F |
| Cap7-Pac1-F | The same as Cap1-Pac1-F |
| Cap8-Pac1-F | The same as Cap1-Pac1-F |
| Cap9-Pac1-F | The same as Cap1-Pac1-F |
| Cap10-Pac1-F | The same as Cap3-Pac1-F |
| Cap11-Pac1-F | The same as Cap3-Pac1-F |
| Cap12-Pac1-F | The same as Cap3-Pac1-F |
| Cap13-Pac1-F | The same as Cap4-Pac1-F |
| Cap1-Nhe-R | TGCCGCTAGCTTACAGGGGACGGGTAAGGTAACGGG |
| Cap3-Nhe-R | TGCCGCTAGCTCACAAGTTTCGTGTGAGATACCGGG |
| Cap4-Nhe-R | TGCCGCTAGCTTACAGGTGGTGGGTGAGGTAGCGGG |
| Cap5-Nhe-R | TGCCGCTAGCTTAAAGGGGTCGGGTAAGGTATCGGG |
| Cap6-Nhe-R | TGCCGCTAGCTTACAGGGGACGGGTGAGGTAACGGG |
| Cap7-Nhe-R | TGCCGCTAGCTTACAGATTACGGGTGAGGTAACGAG |
| Cap8-Nhe-R | TGCCGCTAGCTTACAGATTACGGGTGAGGTAACGGG |
| Cap9-Nhe-R | TGCCGCTAGCTTACAGATTACGAGTCAGGTATCTGG |
| Cap10-Nhe-R | TGCCGCTAGCTTACAGATTACGTGTCAGATAACGAG |
| Cap11-Nhe-R | TGCCGCTAGCTTACAAATGATTAGTCAAATAACGAG |
| Cap12-Nhe-R | TGCCGCTAGCTTACAAGTGGTGGGTGAGGAAACGGG |
| Cap13-Nhe-R | The same as Cap9-Nhe R |
| BamH-GFP-F | TACGGATCCATGGTGAGCAAGGGCGAG |
| EcoR-GFP-R | ATCGAATTCTTACTTGTACAGCTCGTCCATG |
| Q-Bac-F | CCGTAACGGACCT CGTACTT |
| Q-Bac-R | CCGTTGGGATTTGTGGTAAC |
| Q-WPRE-F | CCGTTGTCAGGCAACGTG |
| Q-WPRE-R | AGCTGACAGGTGGTGGCAAT |

Restriction enzyme sites are underlined. F: Forward, R: Reverse.
